# Computational model of brainstem circuit for state-dependent control of hypoglossal motoneurons

**DOI:** 10.1101/199117

**Authors:** Mohsen Naji, Maxim Komarov, Giri P. Krishnan, Atul Malhotra, Frank Powell, Irma Rukhadze, Victor B. Fenik, Maxim Bazhenov

## Abstract

In patients with obstructive sleep apnea (OSA) the pharyngeal muscles become relaxed during sleep, which leads to a partial or complete closure of upper airway. Empirical studies suggest that withdrawal of noradrenergic and serotonergic drives importantly contribute to depression of hypoglossal motoneurons during rapid eye-movement (REM) sleep and, therefore, may contribute to OSA pathophysiology; however, specific cellular and synaptic mechanisms remain unknown. It was recently suggested that, in order to explain experimental observations, the neuronal network for monoaminergic control of excitability of hypoglossal motoneurons has to include excitatory and inhibitory perihypoglossal interneurons that would mediate noradrenergic and serotonergic drives to the motoneurons. In this study, we applied a biophysical network model to validate the rationality of the proposed circuit and to investigate the dynamics of its neuronal populations during REM sleep-induced withdrawal of noradrenergic and serotonergic drives. The state-dependent activity of the model hypoglossal motoneurons during simulated REM sleep with or without a virtual application of noradrenergic and serotonergic drugs was in qualitative agreement with in vivo data. The study predicts the dynamics of the perihypoglossal interneurons during these conditions and corroborates the hypothesis that the excitatory interneurons may integrate both noradrenergic and serotonergic drives. The latter drive has to be mediated by the inhibitory interneurons. The study suggests that perihypoglossal interneurons may serve as novel potential targets for pharmacological treatment of OSA.

## Introduction

Dynamical neuromuscular control of the upper airway dilator muscles differs during wakefulness and sleep; this causes repetitive partial or complete obstruction of upper airway in obstructive sleep apnea (OSA) patients whose upper airways commonly have anatomical features that reduce the size of airway orifice (Horner et al. 1989; Remmers et al. 1978). During wakefulness, a neuromuscular compensation overcomes these anatomical deficiencies and maintains patency of upper airway allowing OSA patients to breath (Mezzanotte et al. 1992). However, this compensation is overpowered by sleep-related depressant mechanisms that cause relaxation of the pharyngeal muscles leading to obstructive events during sleep (Suratt et al. 2014). The upper airway muscle tone and the airway patency is further reduced during rapid eye movement (REM) sleep, which is consistent with the increased severity of obstruction episodes during REM sleep (Eckert et al. 2009; Trinder et al. 2014). The neurochemical nature of neither the neural compensation of pharyngeal muscles during wakefulness nor the depression of these muscles during sleep are well understood.

The activity of hypoglossal motoneurons that innervate tongue and pharyngeal muscles, including the genioglossus (GG), are essential for maintaining upper airway patency, especially during inspiration. Methods including electrical stimulation of hypoglossal nerve (Malhorta 2014; Schwartz et al. 2014; Strollo et al. 2014) as well as chemostimulation (Pillar et al. 2000) and pharmacological interventions (Chan et al. 2006; Fenik et al. 2004, 2005a, 2005b; Fleury Curado et al. 2017; Grace et al. 2013; Horton, et al. 2017; Morrison et al. 2003; Sood et al. 2005; Steenland et al. 2006) have been applied to hypoglossal motoneurons to maintain pharyngeal patency. Understanding of neurological mechanisms of sleep-related changes in neurochemical drive to hypoglossal motoneurons may help developing pharmaceutical treatment for OSA (Jordan et al. 2014).

At present, there is no consensus regarding exact mechanisms responsible for REM sleep-related depression of hypoglossal motoneurons (REM-HD) (Bellingham et al. 1996; Chan et al. 2006; Fenik et al. 2004, 2005a, 2005b, 2015a; Fung and Chase 2015; Grace et al. 2013; Kodama et al. 2003; Kubin et al. 1992, 1993; Lai et al. 2001; Lydic 2008; Morrison, et al. 2003; Parkis et al. 1995; Sood et al. 2005; Steenland et al. 2006; Yamuy et al. 1999). Multiple neurotransmitters have been suggested to play a role in the control of hypoglossal motoneurons during REM sleep: glycine (Fung and Chase 2015; Kodama et al. 2003; Yamuy et al. 1999), GABA (Kodama et al. 2003), serotonin (5-HT) (Fenik et al. 2005b; Lai et al. 2001; Kubin et al. 1992), noradrenaline (NA) (Chan et al. 2006; Fenik et al. 2005b; Lai et al. 2001; Yamuy et al. 1999), glutamate (Bellingham and Berger 1996) and acetylcholine (ACh) (Bellingham and Berger 1996; Grace et al. 2013). However, the most conclusive results were obtained in studies in which application of receptor antagonists that target glycine, GABA_A_, 5-HT and α1-adrenergic receptors into the hypoglossal nucleus abolished REM-HD during carbachol-induced REM sleep-like state in anesthetized rats (Fenik et al. 2004). Follow-up experiments revealed that antagonizing α1-adrenoceptors and 5-HT receptors was necessary and sufficient to abolish REM-HD (Fenik et al. 2005b). Importantly, similar findings were obtained in a series of studies conducted in chronically implanted behaving rats, which were designed to test the role of glycinergic, GABAergic, 5-HT and NA transmission in depression of genioglossus muscle activity during REM sleep in behaving rats (Chan et al. 2006; Morrison et al. 2003; Sood et al. 2005).

Since REM-HD was abolished with a considerable delay of 30-60 minutes following injection of the antagonists into the hypoglossal nucleus (Fenik et al. 2004, 2005a, 2005b), it was proposed that the antagonists had to diffuse outside the hypoglossal nucleus to block receptors, which are located on interneurons mediating the aminergic drive to hypoglossal motoneurons (HM) (Fenik et al. 2005a). Additional analysis of the antagonist effects and their time-courses allowed developing a hypothetical brainstem neuronal network that controls the state-dependent excitability of HM (Fenik 2015a). The key elements of this network were pontine noradrenergic A7 and medullary raphe 5-HT neurons that are REM-OFF neurons (Fenik et al. 2015b; Heym et al. 1982; Rukhadze et al. 2008; Trulson and Jacobs 1979); and the hypothetical excitatory and inhibitory interneurons, that are likely to be located in perihypoglossal region and mediate noradrenergic and 5-HT drives to HMs (Fenik 2015a). In this study, we developed a computational network model to test this hypothesis and to investigate the impact of withdrawal of noradrenergic and serotonergic drives during the transition from non-REM (NREM) sleep to REM sleep on HM activity.

## Materials and methods

### A. Biophysical model

Since little is known about intrinsic properties of HMs, we assumed that activity of HMs is linearly related to the excitatory drive from excitatory perihypoglossal interneurons (EPI). To model activity of EPI, we considered a network model that contained four neuronal populations: A7 noradrenergic neurons, 5-HT neurons in raphe nuclei, inhibitory perihypoglossal interneurons (IPI; which are likely to be GABAergic) and EPI, connected to each other as shown in Figure 2A.

### Models for individual neuron dynamics

The populations of the IPI and EPI were modeled using Hodgkin-Huxley formalism. The membrane potential of each neuron was governed by the following equation (Kilpatrick and Ermentrout 2011; Komarov and Bazhenov 2016; Traub 1982):

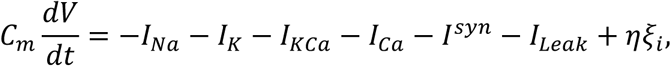

where *V* represents transmembrane voltage of the neuron, *C*_*m*_ stands for membrane capacitance, *I*_*na*_ is fast Na^+^ current, *I*_*k*_ is a delayed rectifier K^+^ current, *I*_*kca*_ is slow Ca^2+^ dependent K^+^ current and *I*_*ca*_ is a high-threshold activated Ca^2+^ current, *I*_*leak*_ is a leak current and the term η ξ_*i*_*(t)* corresponds to the fluctuations in the transmembrane current (representing spontaneous background activity) which are given by a white noise process with the following properties: *<*ξ_*i*_ *(t) >= 0, <* ξ_*i*_ *(t)* ξ_*i*_ *(t – t*_*0*_)_=_ >= δ *(t – t*_*0*_). Each ionic current was modeled using Hodgkin-Huxley formalism:

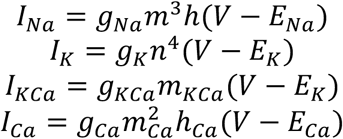

where *g*_*x*_ is a maximal conductance, *E*_*x*_ is a reversal potential and *m*_*x*_ and *n*_*x*_ are activation and inactivation gating variables correspondingly (subscript x denotes one of the ionic currents *x* ∈{*Na, K, KCa, Ca*}). The gating variables obey the following equation:

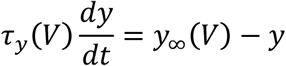

where y is one of the gating variables, and voltage dependent functiony_∞_(*V*) and τ (*V*) are:

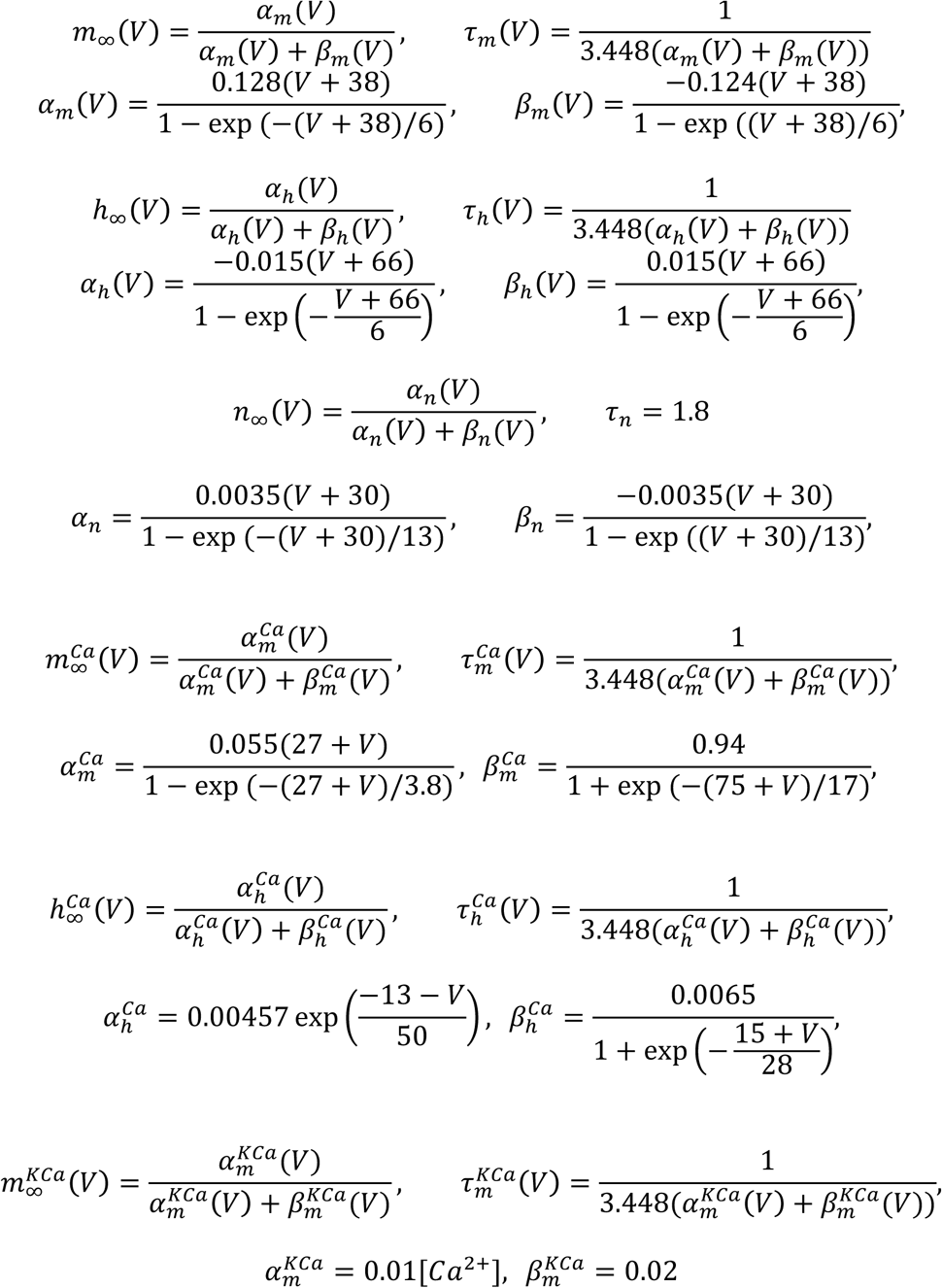

Dynamics of calcium concentration [*Ca*^*2+*^] obeys the following equation:

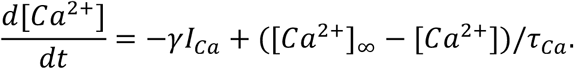

The leak current consists of two parts: *I*_*leak*_ *= I*_*k*_ *+ I*_*cl*_ where *I*_*cl*_ *= g*_*cl*_*(V-V*_*CL*_) is a chloride leak current and 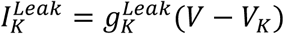 is a potassium leak current.

Spiking activity of populations of A7 and RN neurons were modeled using Poisson process, which characteristic frequency depended on the sleep stage.

### Models for action of neuromodulators (5-HT and NA) and drugs (Methysergide and Prazosin)

#### Interneurons

The scheme presented in Figure 2A assumes that each neuron in the population of IPI contains serotonergic 5-HT receptors, which are able to produce cell inhibition through opening of K^+^ channels. The dynamics of 5-HT receptors under action of serotonin (5-HT) and Methysergide (Me) was modeled based on the following transition scheme (Destexhe et al. 1998):

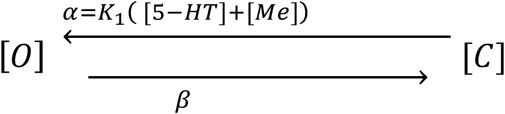

here labels [O] and [C] denote opened and closed state of the 5-HT receptor correspondingly; α and *β* denote rates of transition between the states. The rate of transition from closed to open state is a linear function of the sum of 5-HT and Me concentrations α=*K_1_*([*5-HT*]+[*Me*]) The rate of transition from opened to the closed state is constant (β). The described above scheme leads to the following first-order kinetic equation for the fraction of the opened serotonergic receptors:

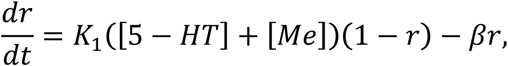

which, in turn, produce activation of intracellular G-protein:

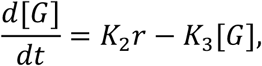

Finally, activation of G-protein modulates potassium leak current in the following way:

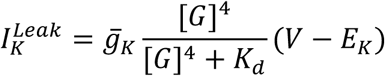

The scheme presented in Figure 2B assumes that each neuron in population of EPI interneurons contains α1-adrenergic receptors, whose activation can facilitate cell activation. The dynamics of NA receptors under action of noradrenaline and Prazosin (Pz) was modeled based on the following transition scheme:

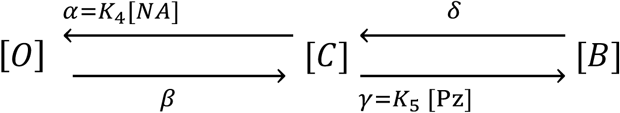

here [O], [C] and [B] denote opened, closed and blocked state correspondingly. NA and Pz compete for the noradrenergic receptors, transforming them to either opened (NA) or blocked (Pz) states. The rate of transition of closed to opened state depends on the concentration of NA: α=*K*_4_[*NA*], while transition from closed to blocked state is controlled by concentration of Pz: γ = *K*_5_[*PZ*] *K*_*4,5*_ are positive constants. The described above scheme leads to the following equations for fraction of opened, closed and blocked receptors:

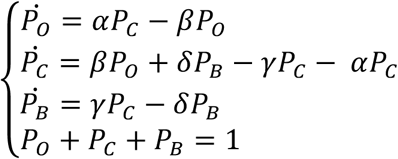

Similar to the case of IPI, opening of NA receptors leads to activation of the second messenger and modulation of K^+^ currents:

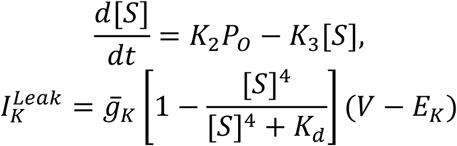

#### Parameters

Parameters for IPI: *C*_*m*_*=1* μ *F/cm*^*2*^, *E*_*Na*_*=55 mV*, *E*_*+*_*=-85 mV:E*_*cl*_*=-71Mv,E*_*ca*_*=120mV, g*_*na*_*=150 ms/cm*^2^, *g*_*k*_ *2009= 10 ms/cm*^*2*^*, g*_*kca*_ *= g*_*ca*_ *= 0 ms/cm*^*2*^*,g*_*cl*_*= 0.015 ms/cm*^*2*^, τ_*ca*_ *= 400 ms, [Ca*^*2+*^]∞ *= 2.4 · 10*^*-4*^.

Parameters for EPI: *C*_*m*_= μ*F/cm*^*2*^, *E*_*Na*_*=55 mV*, *E*_*K*_ *= -85 mV, E*_*CL*_ *= -71 mV, Eca=120mV, g*_*Na*_ *= 75 ms/cm*^*2*^, *g*_*+*_ *= 25 ms/cm*^*2*^*, g*_*kca*_*= 1.5 ms/cm*^*2*^*, g*_*ca*_ *= 0.2 ms/cm*^*2*^*,g*_*cl*_ *= 0.022 ms/cm*^*2*^, τ_*ca*_*= 400 ms, [Ca*^*2+*^]∞ *= 2.4 · 10*^*-4*^.

## Results

### Pharmacological interventions reveal interactions between brainstem nuclei in the hypoglossal motoneurons control

A conceptual model of a brainstem neural network that can potentially explain in vivo data on state-dependent control of hypoglossal motoneurons has been proposed in (Fenik 2015a). In this model, the main neuronal populations that are responsible for the state-dependent control of HMs include five distinct populations of cells (Figure 1A, from (Fenik et al 2015b)): 5-HT raphe neurons (RN), noradrenergic A7 neurons, local inhibitory possibly GABAergic interneurons, excitatory interneurons (RF-neurons), and hypoglossal motoneurons (HMs). HMs are indirectly controlled by A7 neurons through the populations of RF-neurons. The RN provided inhibitory 5-HT projections to the inhibitory interneurons that, in turn, inhibit the RF-neurons. In vivo data behind this conceptual model are summarized below.

Recordings from 5-HT neurons within nucleus raphe pallidus showed the highest activity during wakefulness (mean: 4.85±0.37 spikes/s) with gradually slowed activity during NREM sleep (mean: 3.76±0.36 spikes/s) and the least activity during REM sleep (mean: 0.92±0.23 spikes/s) (Heym et al. 1982). We also obtained preliminary data that noradrenergic A7 neurons have tonic low-frequency discharges during NREM sleep (mean: 0.97±0.4 spikes/s) whereas they are almost silent (mean: 0.092±0.09 spikes/s) during REM sleep (Fenik et al. 2015b). Figure 1B summarizes mean activity of A7 and raphe pallidus neurons during NREM and REM sleep.

The injection of a solution of prazosin (Pz) and methysergide (Me) into the hypoglossal nucleus abolished REM sleep-related depression of HMs (REM-HD) that was induced by pontine injection of carbachol in anaesthetized rats (Fenik et al. 2005b). These findings are summarized in Figure 1C (modified from (Fenik et al. 2005b)). Following 27–83 min after the combined antagonist injections, hypoglossal nerve activity was disfacilitated to 27.0% of baseline, approximately the level measured during control carbachol responses, and carbachol injected at this time period was unable to further reduce activity in hypoglossal nerve.

Two other separated trials with injections of “prazosin only” and “methysergide only” were conducted to quantify the contribution of noradrenergic and serotonergic neurotransmission to REM-HD. Following the “prazosin only” injections, the HM activity was reduced to 20.8% of baseline, below the control carbachol level (26.1%), and further reduced by another carbachol injection to 14.8%. Thus, despite the fact that HM activity was reduced by prazosin more than by the mixture of prazosin and methysergide, the reduced activities before and after carbachol injection had a significant difference revealing a remaining REM-HD of about 6%. Using these numbers, we can calculate relative contribution of 5-HT and NA mechanisms to REM-HD as following: the estimated relative contribution of serotonergic mechanisms to REM-HD is (20.8-14.8)/(100-14.8)=7%, and the relative contribution of noradrenergic mechanism is (100-20.8)/(100-14.8)=93% (see also (Fenik 2015)).

After the “methysergide only” injections, the HM activity was reduced only to 85.8% of baseline and carbachol injections elicited a strong REM-HD. Note that following methysergide the HM activity after carbachol administration was significantly higher (37.6%) than during the control REM sleep-like state before the antagonists (27.2% on baseline activity) suggesting a disinhibitory effect of methysergide on REM-HD (Fenik et al. 2005b).

Collectively, these data suggest that prazosin may be fully responsible for the disfacilitation, whereas methysergide, for disinhibitory effect on REM-HD.

**Figure 1.**
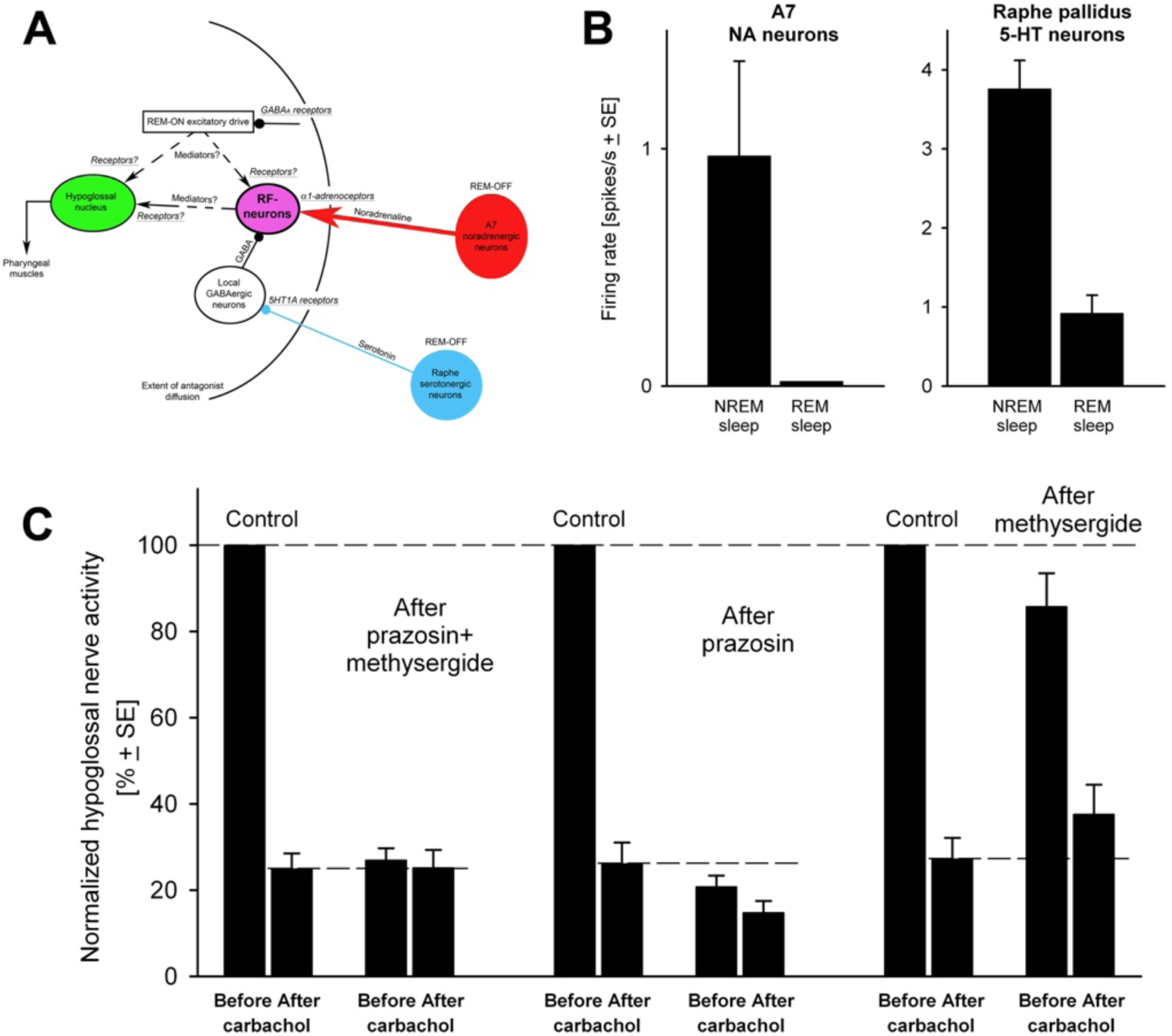
Empirical data that were used to construct and validate the biophysical model of a network to provide monoaminergic control of HM activity. A) A brainstem circuit that illustrates the main neural populations responsible for state-dependent control of HM (from (*Fenik 2015a*)). B) Recordings from A7 (*Fenik et al. 2015b*) and RN (*Heym et al. 1982*) revealed that they have lower activity during REM compared to NREM sleep. C) Hypoglossal nerve activities during carbachol-induced REM sleep-like episodes that were evoked before and after the injection of methysergide, prazosin or a mixture of prazosin and methysergide.

Based on the described above physiological data, we designed a biophysical model representing dynamics of the main neuronal groups within the network responsible for the state-dependent control of HM and predicted changes in their behavior under different physiological conditions, such as during NREM and REM sleep, and under different concentrations of Pz and Me.

### Biophysical model

We designed a biophysical model to explore the role of proposed neuronal populations and neurotransmitters in state-dependent control of HMs (Figure 2A). We assumed that the activity of HMs was linearly related to that of the excitatory perihypoglossal interneurons (EPI). This excitatory drive from EPI neurons to HMs depended on the interactions between four distinct neuronal populations: A7 neurons, 5-HT RN, EPI and inhibitory perihypoglossal interneurons (IPI).

#### The baseline model

We adjusted firing rates of A7 neurons and RN in the model to be equal to those obtained in *in vivo* experimental recordings during NREM and REM sleep (Figure 2B,C). The reduction of the firing rate of REM-OFF A7 neurons and RN during the transition from NREM to REM sleep decreases the release of NA and 5-HT neurotransmitters.

The status of α1-adrenoceptors in EPI and 5-HT1A receptors in IPI determined the state of potassium leak currents in these neurons. We assumed that the EPI also received GABAergic inhibitory drive from the IPI that affected the postsynaptic currents of the EPI. Figure 2D,E shows the activity of the IPI and EPI populations during NREM and REM sleep stages in the model. Relatively high activity of RN during NREM sleep inhibited activity of IPI as compared to REM sleep (Figure 2D). The net effect of reduced excitatory drive from A7 neurons and increased inhibitory drive from IPI resulted in a markedly lower activity of EPI during REM sleep as compared to NREM sleep (Figure 2E-F), which was in line with experimental recordings from HMs (Figure 1C). Thus, the behavior of EPI in the baseline model was REM-OFF and mimicked the activity of HMs during NREM and REM sleep whereas the behavior of IPI had the REM-ON pattern, i.e., they were more active during REM sleep as compared to NREM sleep.

**Figure 2.**
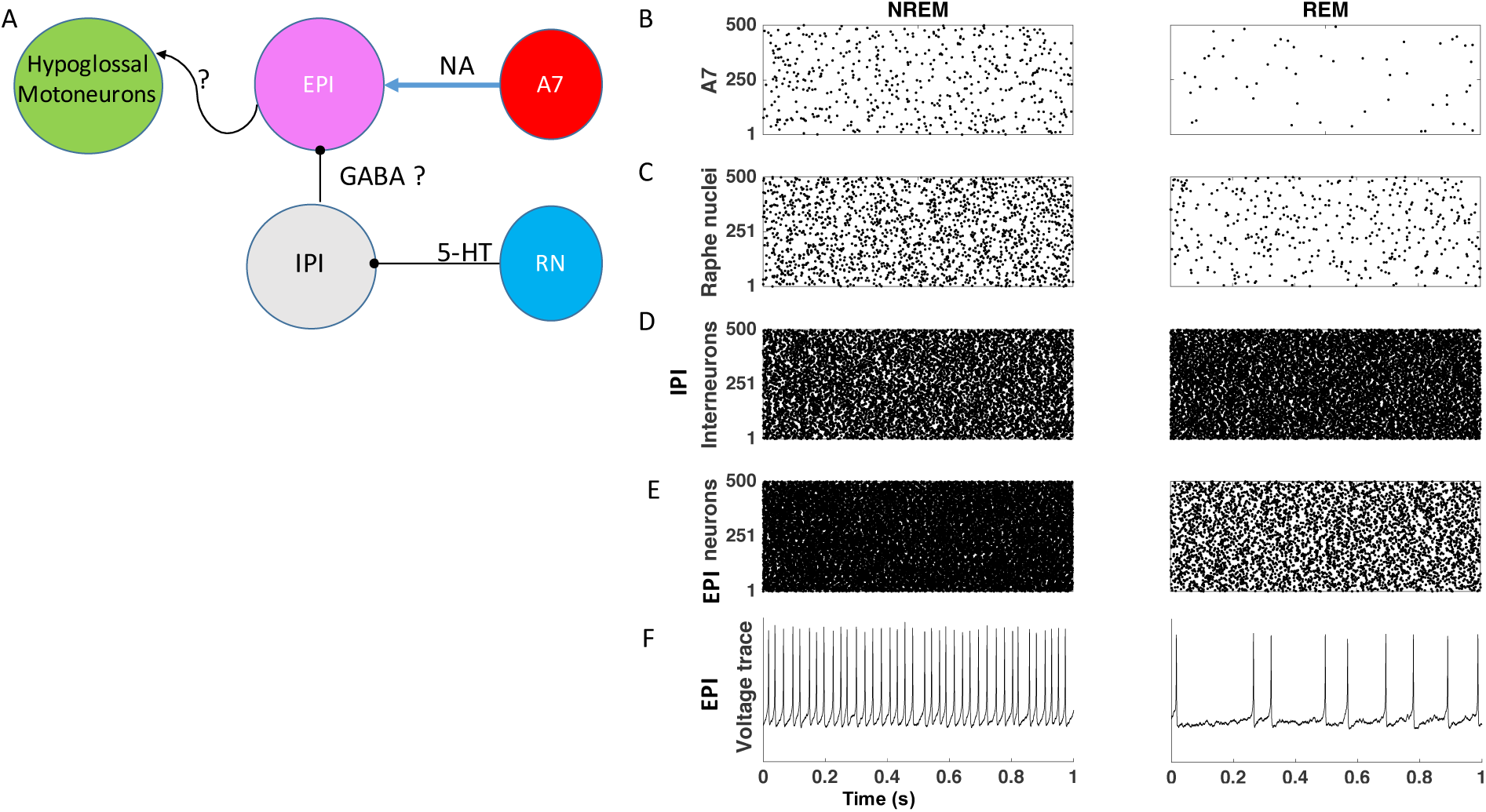
Baseline model activity during REM and NREM sleep. A) Schematic of a biophysical model of brainstem circuit. We assumed that RN inhibit IPI through 5-HT1A receptors and IPI, in turn, inhibit EPI by affecting GABA_A_ receptors. EPI neurons integrate both the excitatory noradrenergic drive from A7 nuclei and inhibitory drive from IPI. The neurotransmitter by which EPI population excite HMs has yet to be identified. B-C) The rasterograms show the A7 and RN firing during NREM and REM sleep. D) The IPI neurons are disinhibited during REM compared to NREM sleep. E-F) The rastrograms and voltage traces show higher activity of EPI during NREM compared to REM sleep.

#### Effects of prazosin and methysergide on the neural behavior in the basic model

Starting from the baseline model, we studied the effect of different extracellular concentrations of Pz, a α1-adrenergic antagonist, and Me. Me is a broad-spectrum 5-HT antagonist. In our model, it worked as an agonist of inhibitory 5-HT1A receptors as proposed previously (Trulson and Jacobs 1979; Scrogin et al. 2000). The levels of the concentration of Pz and NA in the model varied within a broad range of values that competed to affect the probability of α1-adrenoceptors located on the EPI neurons to be in the open or closed states (Figure 3A,C). An increase in the Pz level and a decrease of the NA drive resulted in the reduction of the probability of α1-adrenoceptors to be open (Figure 3C). Conversely, Me worked as agonist and helped 5-HT to increase the probability of the 5-HT1A serotonergic receptor of the IPI to be in the open state (Figure 3B, D).

**Figure 3.**
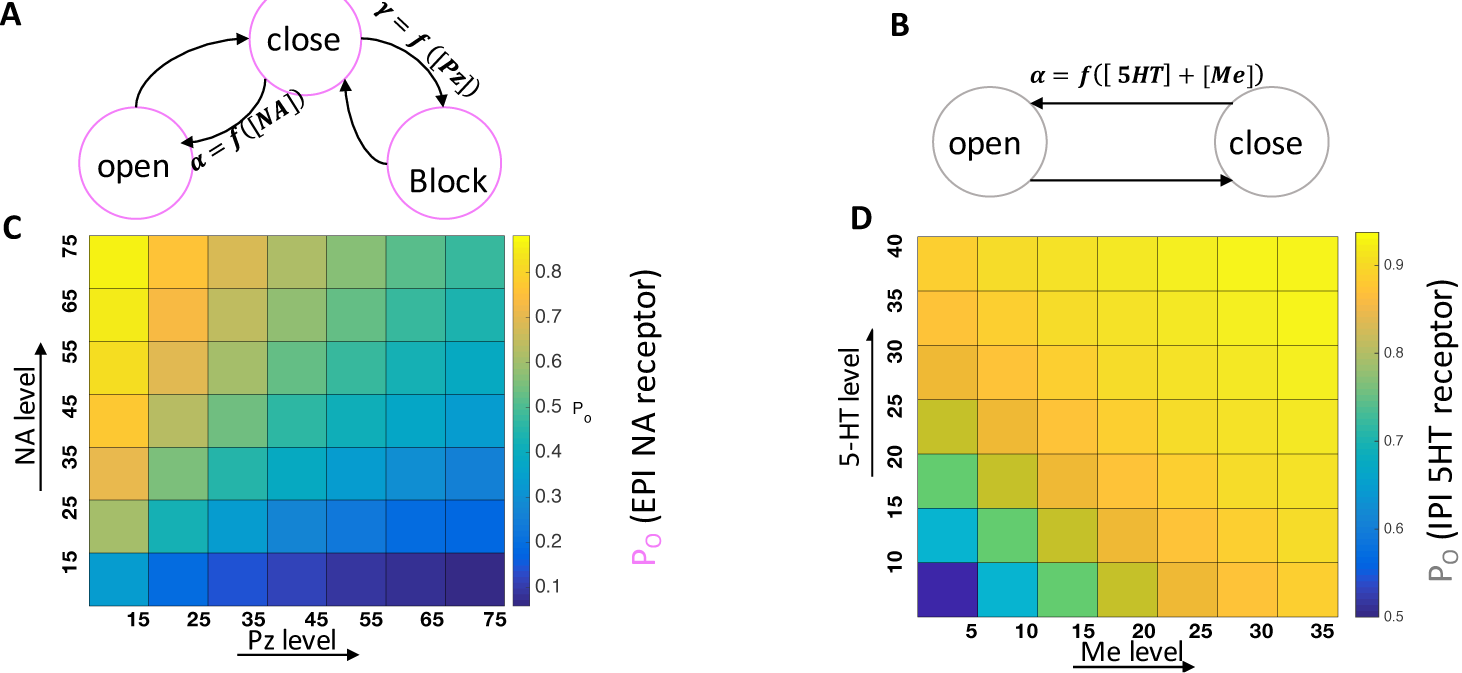
The effect of different ligand concentrations on the state of α1-adrenergic and 5-HT1A receptors. A) The dynamics of NA receptor under action of NA and Pz was modeled based on the transition between open, close and blocked states (see Methods). B) The dynamics of 5-HT1A receptor under the action of 5-HT and Me was modeled based on the transition between open and close states. C) Pz competed with NA to reduce the probability of the NA receptor to be in the open state. D) Me helped 5-HT to increase the probability of the 5-HT1A receptor to be in the open state.

Figures 4A and 4B illustrate the effect of different levels of Pz and Me on the firing rate of EPI during NREM and REM sleep. In both cases, increasing the prazosin level decreased the EPI activity, while increasing methysergide level slightly increased the EPI activity. Effect of the prazosin was most prominent during NREM sleep, which is illustrated by plotting EPI firing vs Pz level (see inset in Figure 4A). Thus, the model explained the effect of competition between prazosin and methysergide in controlling activity of the EPI population. Next, we selected the values for Pz and Me to match model responses to the empirical data of the agonists application across sleep stages (Figure 1). Existence of the match supports our hypothesis that proposed network circuit (Figure 2A) is sufficient to explain EPI (and therefore HM) behavior under control conditions and following drug applications.

The results of the model behavior are summarized in Figure 4C-F. The antagonists, Pz and Me, affected receptors that are located on EPI and IPI neurons, respectively. Therefore, they changed the activity of IPI and EPI populations but not the firing rates of A7 or RN in our network model (Figure 4C-F). When the mixture of Pz and Me was applied, the EPI activity during NREM sleep was reduced to 24.59% of the baseline (the activity during NREM sleep under a drug-free condition). This level was similar to the level of the EPI activity during REM sleep in control (24.27% of the baseline). During the following REM sleep, the activity of EPI minimally decreased to 21.97% of baseline, which mimicked the experimentally observed abolition of REM-HD by Pz+Me.

In the “prazosin only” condition, the EPI activity reduced to 24.18% of baseline during NREM sleep and further reduced to 20.23% during REM sleep. This level of EPI activity was significantly (p<0.05, n=10; paired t-test) less compared to that during REM sleep after Pz+Me (21.97%) suggesting that Me has a disfacilitatory effect on EPI activity, which was also observed in *in vivo* experiments (Kodama et al. 2003). The estimated relative contribution of serotonergic mechanisms to REM-HD in the model was (24.18-20.23)/(100-20.23) ≈5%, and the relative contribution of noradrenergic mechanisms was 95%, which is consistent with the *in vivo* findings. Thus, our biophysical model confirmed that noradrenergic disfacilitation during REM sleep is the major mechanism that is responsible for depression of HM.

Under the “methysergide only” condition, the activity of IPI population was reduced during NREM sleep as compared to baseline, which resulted in disinhibition of the EPI population to 101.3% of baseline (Figure 4E,F). During the following REM sleep, the activity of IPI population did not change because Me occupied all 5-HT1A receptors, which made these neurons insensitive to state-dependent changes in 5-HT level (Figure 4E). The activity of EPI group was largely depressed during REM sleep (Figure 4F). However, it remained significantly higher than that during control REM sleep (p<0.05, n=10; paired t-test) mimicking the disinhibitory effect of methysergide on the REM-HD observed *in vivo* (see Figure 1C).

To summarize, the model fully mirrored the behavior of HMs that were observed *in vivo*. The only discrepancy was the minor increase of activity of EPI that occurred during NREM sleep after application of Me in contrast with *in vivo* data that show the reduced activity of HMs during NREM sleep under the “methysergide only” condition. (see Figure 1C). The reason for this discrepancy can be explained by the fact that in our model we did not include the early direct effect of Me on HMs, which decreased activity of HMs immediately after Me injections but did not affect REM-HD *in vivo* (Fenik et al. 2005b). Taking this caveat into account, we concluded that the model confirms that serotonergic drive may be sufficient to explain disinhibition of HMs during REM sleep.

**Figure 4.**
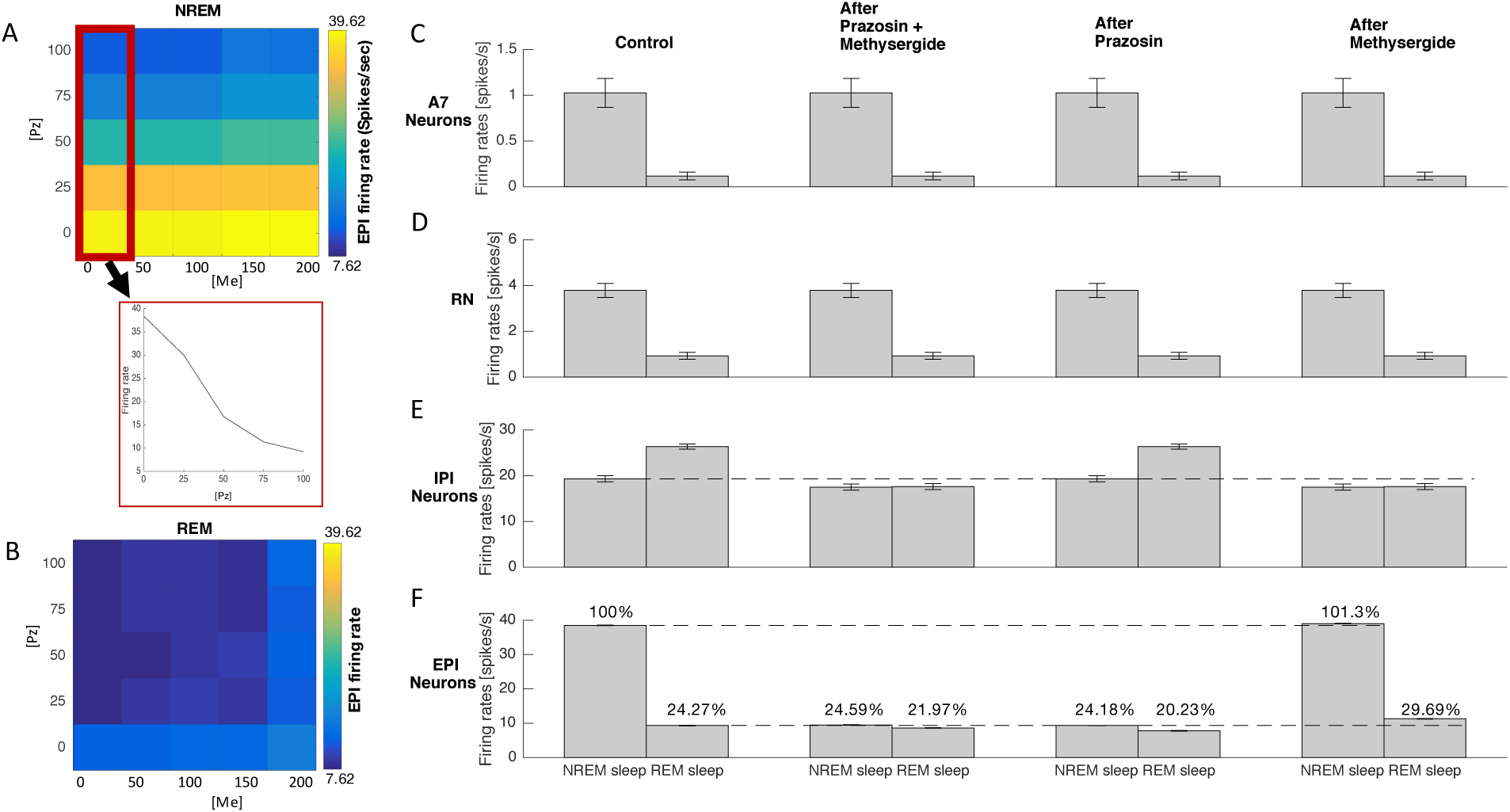
Simulated effects of drug application in the model. A) Effect of different levels of Pz and Me on the firing rate of EPI during NREM sleep. The inset shows, for the minimum level of Me, that increasing the Pz level reduced the activity of EPI. B) Effect of different levels of Pz and Me on the firing rate of EPI during REM sleep. C-D) The firing rates of A7 and RN during NREM and REM sleep, which were independent of the presence of either antagonist. E) IPI neurons were disinhibited during REM sleep in control. The application of Me inhibited their firing rate below the control NREM sleep level and made them state-independent, i.e. insensitive to changes of the level of 5-HT. F) The firing rate of EPI during NREM and REM sleep under the drug-free control condition and after application of antagonists.

## Discussion

*In vivo* data suggest complex interaction between different types of neurotransmitters in the control of HMs activity during sleep. Different brainstem circuits may be involved and understanding of specific mechanisms is lacking, which negatively impacts developing novel therapeutic approaches. The goal of this new study was to construct a biophysical model of the brainstem neuronal network that may explain state-dependent activity of HMs and to test this model against *in vivo* data, including transitions from NREM to REM sleep and application of noradrenergic and serotonergic antagonists, to either support or refute the proposed hypothetical minimal network of HMs modulation and control.

Excitatory and inhibitory perihypoglossal interneurons have been suggested to mediate state-dependent 5-HT and NA drives to HMs (Fenik 2015a). Our network model included four distinct interacting neuronal groups: A7 noradrenergic neurons, serotonergic raphe neurons (RN), excitatory perihypoglossal interneurons (EPI), and inhibitory perihypoglossal interneurons (IPI), populations. In the model, REM-HD occurred when the A7 neurons and RN fired at their minimum rate. The behavior of EPI neurons in the model corroborates that the state-dependent control of HM excitability could be mediated by EPI neurons that integrate both the excitatory noradrenergic drive from A7 nuclei and disinhibitory serotonergic drive from RN, which is mediated by IPI. Manipulating with the status of α1-adrenoceptors in EPI and 5-HT1A receptors in IPI groups confirmed (1) that noradrenergic disfacilitation and serotonergic disinhibition can be responsible for REM-HD; and (2) that the noradrenergic mechanisms are the major contributor to hypoglossal depression during REM sleep-like state as compared to the contribution of 5-HT mechanisms.

We constructed the model based on experimentally obtained results that combined injections of α1-adrenergic and 5-HT receptor antagonists into the hypoglossal nucleus abolished REM-HD during carbachol-induced REM sleep-like state in anesthetized rats (Fenik et al. 2005b). The NA drive proved to be mostly responsible for REM-HD as compared to 5-HT drive under those experimental conditions (Fenik 2015a). These findings were confirmed by Richard Horner group in chronically implanted behaving rats with recording of genioglossus (GG) muscle activity during natural REM sleep (Chan et al. 2006). The application of terazosin, a α1-adrenoceptor antagonist, into the hypoglossal nucleus via microdialysis probe significantly decreased REM-HD in respiratory-modulated activity of GG by approximately 50% compared to saline controls (Chan et al. 2006). This decrease of REM-HD was much smaller than the approximately 90% decrease of REM-HD that was observed in anesthetized rats after injections of prazosin (Fenik et al. 2005b, 2015a), which may have at least two explanations. First, the concentration of terazosin that entered the hypoglossal nucleus from the microdialysis probe in this study (Chan et al. 2006) is unknown. Thus, following diffusion, its concentration at the vicinity of EPI could be insufficient to block all relevant α1-receptors; therefore, only partial effect could be observed in that study. The dose-response experiments were not reported to see if the effect of terazosin was saturated. In addition, we showed (see inset in Figure 4A) that increasing Pz concentration in the model could reduce the activity of EPI neurons. So, it supports the suggestion that the smaller effect of terazosin could be related to smaller drug concentrations. Second, the relatively small effect of terazosin may imply that, in behaving rats, additional neurotransmitter mechanisms may contribute to REM-HD during natural REM sleep, e.g., cholinergic inhibition (Grace et al. 2013). However, the magnitude of terazosin-induced decrease of REM-HD appeared to be the largest among the effects on REM-HD that were produced by the application of antagonists of glutamatergic (Steenland et al. 2006), serotonergic (Sood et al. 2005), GABAergic (Morrison et al. 2003), glycinergic (Morrison et al. 2003) or muscarinic receptors (Grace et al. 2013) using the same approach under the same experimental conditions.

In addition, phenylephrine, a α1-adrenergic agonist, was applied in attempt to rescue activity of HM during REM-HD in behaving rats (Chan et al. 2006). The application of phenylephrine increased respiratory-modulated GG muscle activity proportionally to that during control saline application in each behavioral state: wakefulness, NREM and REM sleep. Thus, there was no apparent change in REM-HD following the phenylephrine application suggesting that the agonist could not reduce REM-HD in behaving animal. The failure of phenylephrine to rescue GG activity during REM sleep is readily explained by the following. Again, the amount of phenylephrine that leaves the probe and its actual concentration within the hypoglossal nucleus is unknown but it can be assumed to be similar to that of terazosin (the concentration of both drugs was 1 mM in the probe) (Chan et al. 2006). According to our experience, microinjections of phenylephrine into the hypoglossal nucleus in anaesthetized rats produce relatively short-lasting responses (approximately 15 min) (Fenik et al. 1999) as compared to the effect of prazosin, which lasted more than 3 hours (Fenik et al. 2005b). This suggests that the removal of phenylephrine from extracellular fluid occurs at a larger rate than that of prazosin. Since chemical formulas of prazosin and terazosin are almost identical, their pharmacokinetics is likely to be similar. Thus, probably larger concentration of phenylephrine needs to be applied into the hypoglossal nucleus in order to reach EPI by diffusion. Based on this reasoning, it most likely that phenylephrine did not reach EPI in behaving rats where we believe it could rescue the activity of HMs during REM-HD. However, the moderate direct effect of phenylephrine on HMs in behaving rats is consistent with our experience in anesthetized rats (Fenik et al. 1999).

In the model proposed in our new study, NA drive affects HMs indirectly, via EPI that have a net excitatory effect on HMs. The need for this group of interneurons was dictated by the fact that α1-adrenergic and 5-HT receptor antagonists injected into HMs required 30-60 min to reach their target receptors in the carbachol model of REM sleep (Fenik et al. 2004, 2005a, 2005b). Recently, we obtained preliminary data that support the hypothesis that NA drive to HMs is not direct (Fenik and Rukhadze 2016). The question which neurotransmitter is used by EPI to excite HMs remains open. One possible candidate is glutamate, which is the most widely used excitatory neurotransmitter in the central nervous system. However, in experiments in chronically implanted behaving rats, effects of glutamate antagonists on REM-HD were unexpectedly weak (Steenland et al. 2006). Thus, the neurotransmitters that are responsible for the net excitatory drive from EPI to HMs needs to be identified in future studies.

We proposed that RN inhibit IPI through 5-HT1A receptors and IPI, in turn, inhibit EPI by GABA_A_ receptors. More experiments are needed to confirm that GABA_A_ and 5-HT1A receptors in the brainstem circuit are involved in the state-dependent control of HM activity. Nevertheless, our model provides evidence that interneurons, such as EPI and IPI that mediate effects of A7 neurons and RN on HMs, may be necessary to explain the dynamics of HMs during different sleep stages and under different drug conditions. Indeed, a simple model with direct inputs from A7 neurons and RN to HM fails to explain their complex behavior. We believe that the model proposed in our new study can be enriched with emerging data from *in vivo* experiments and can be used for further understanding the HMs dynamics in the normal and the pathological states.

To conclude, we developed a computational network model to investigate the impact of withdrawal of noradrenergic and serotonergic drives during the transition from NREM to REM sleep on HM activity. This model dynamics are consistent with a broad range of empirical data and it makes predictions about specific minimal circuit interactions that are sufficient to explain the observed in vivo phenomena. While we cannot exclude that the other neuronal circuits may be also involved, the results of our study predict the dynamics of the excitatory and inhibitory perihypoglossal interneurons, which is consistent with in vivo recordings, during different sleep stages and under various pharmacological manipulations. Importantly, our study suggests new targets for clinical interventions that may reduce severity of obstruction events in OSA patients.

## Acknowledgements

This work was supported by grants from ONR (MURI: N000141612829), National Heart, Lung, and Blood Institute (HL116845), and NIH (RO1 HL081823, RO1 HL085188, K24 HL132105, and T32 HL134632).

